# Autoencoders with shared and specific embeddings for multi-omics data integration

**DOI:** 10.1101/2024.08.14.607979

**Authors:** Chao Wang, Michael J. O’Connell

## Abstract

**Motivation:** In cancer research, different levels of high-dimensional data are often collected for the same subjects. Effective integration of these data by considering the shared and specific information from each data source can help us better understand different types of cancer.

**Results:** In this study we propose a novel autoencoder (AE) structure with explicitly defined orthogonal loss between the shared and specific embeddings to integrate different data sources. We compare our model with previously proposed AE structures based on simulated data and real cancer data from The Cancer Genome Atlas. Using simulations with different proportions of differentially expressed genes, we compare the performance of AE methods for subsequent classification tasks. We also compare the model performance with a commonly used dimension reduction method, joint and individual variance explained. In terms of reconstruction loss, our proposed AE models with orthogonal constraints have a slightly better reconstruction loss. All AE models achieve higher classification accuracy than the original features, demonstrating the usefulness of the embeddings extracted by the model. Particularly, we show that the proposed models have consistently high classification accuracy on both training and testing sets. In comparison, the recently proposed MOCSS model that imposes an orthogonality penalty in the post-processing step has lower classification accuracy that is on par with JIVE.

**Availability and Implementatio:** The relevant datasets and models developed in this study are available in the following GitHub repository: https://github.com/wangc90/AE_Data_Integration.

## Introduction

Data integration is the process of combining data from different modalities or sources that each provide a separate view of the common phenomenon. Compared with the analysis of the individual data modality, different data sources can provide complementary information thus promising a more comprehensive understanding of the system [14]. This is particularly important in the biomedical field, where multi-omics data including genomics, transcriptomics, proteomics, epigenomics, metabolomics, and meta-genomics are increasingly made available for different study objectives. One of the most important examples is The Cancer Genome Atlas (TCGA), a project that has collected different types of omics data for over 20,000 tumor and normal samples, which collectively cover 33 different types of cancers with clinical information for a subset of subjects [13]. In addition to an effective integration strategy to account for the distributional difference of the data source, each modality of data is often also high dimensional. For example, gene expression profiles for each subject in the TCGA database consist of 60,483 measurements (FPKM: fragments per kilobase of transcript per million mapped reads) corresponding to the expression measurement for each gene at the transcript level. DNA methylation profile for each subject was determined by Infinium HumanMethylation 450 BeadChip Arrays containing 485,578 probes that measure the methylation level for a small part of the human genome [16]. The high dimensionality can pose significant analysis challenges for classic statistical methods and machine-learning techniques since the number of features in each modality is larger than the number of observations (*P* ≫ *N*). Consequently, dimension reduction such as principal component analysis (PCA) or customized filtering (*e*.*g*., only considering differentially expressed genes or methylation probes in the analysis) are often needed either before or during the data integration process.

The majority of the data integration methods can be grouped into four different categories: dimension reduction, kernel, network, and deep learning methods [1]. Among these, joint dimension reduction methods are probably the most heavily explored, with integrative extensions of commonly used dimension reduction methods like principal component analysis [4, 3], canonical correlation analysis [17], and non-negative matrix factorization [15].

However, there has recently been more interest in deep learning methods for data integration. Deep learning methods are more flexible, and they can identify non-linear patterns much more easily than most dimension reduction methods, due to the non-linear activation functions of most deep learning methods. Deep learning models can learn the hierarchical representation of data automatically without the constraints of linear relationships, making them particularly suitable and flexible for the integration of multi-omics data in biomedical applications [14].

An autoencoder (AE) is a deep-learning approach to find the latent representation of the input with a lower dimension that contains the necessary information to reconstruct the original input. An AE consists of an encoder function *f* (·) parameterized by *θ* and a decoder function *g*(·) parameterized by *ϕ* such that for a single input **x**, *g*_*ϕ*_(*f*_*θ*_(**x**)) ≈ **x** and the *f*_*θ*_(**x**) is the embedding of the original input and **x**^′^ = *g*_*ϕ*_(*f*_*θ*_(**x**)) is the reconstructed input. The goal of the AE is to capture the essential information in the data by minimizing the reconstruction error. Different metrics can be used to measure the error such as mean squared error: 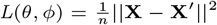,where **X** is the entire dataset and **X**^′^ is the corresponding reconstruction of the entire dataset after applying the AE model [12]. If *f*_*θ*_(·) and *g*_*ϕ*_(·) are linear functions then **X**^′^ will lie in the principle component subspace making the AE similar to PCA. If *f*_*θ*_(·) and *g*_*ϕ*_(·) are nonlinear, the input will then be mapped onto a lower dimensional manifold that is more informative than principle components when there are non-linear interactions in the data [14]. In practice, AEs are neural network architecture that often consists of fully connected layers of encoder and decoder. The encoder takes the input data and compresses them into lower dimensional representations and the decoder takes this lower dimensional representation and reconstructs the input at the final output layer.

A number of different AE architectures with variational components were proposed for multi-omics data integration previously [12]. The most straightforward and easy-to-implement one is to simply concatenate the scaled data source as the input for AE, also referred to as the Autoencoder with Concatenated Inputs (CNC AE) [12]. For this AE architecture, the encoder consists of a number of fully connected layers including the embedding layer to transform the concatenated input of different data sources (*i*.*e*., concatenation of X1, and X2) from a higher dimension to a lower dimensional representation. The corresponding decoder then takes this lower dimensional representation as the input aiming to reconstruct the original input for different data sources separately. A slightly modified version of this CNC AE referred to as X-shaped Autoencoder (X AE), in which the individual data sources are first preprocessed separately before being joined together to be further processed by the model. Finally, instead of directly concatenating the preprocessed data source, a pair-wise mutual concatenation of the input is used in Mixed-Modal Autoencoder (MM AE) model to take advantage of the potentially shared information among the different data sources as suggested in the previous study [12]. Despite these different architectures, all these AE structures fail to take advantage of the potentially joint and individual components/information across the data source in their architectural design.

A recent study implemented the notion of joint and individual components in AE for multi-omics data integration with the multi-omics data clustering and cancer subtyping via shared and specific representation learning (MOCSS) model [2]. Specifically, two separate AEs were created for each data source and designated as the shared and specific respectively. With 2 different data sources, all 4 AE have the same architecture. However, instead of using the concatenated input from different data sources to extract the potentially shared information, 2 separate shared embedding vectors were extracted from each data source based on the aforementioned shared AEs. To encourage these embeddings to be shared from data source to data source, these vectors were first transformed through another fully connected neural network (MLP: multiple linear perceptrons) before a contrastive learning process was used to align them by reducing the pair-wise contrastive loss between these vectors. The orthogonal penalties were applied between the shared embedding vector and the specific embedding vector for each data source individually and then added together as the final orthogonal loss. Despite taking advantage of the potentially shared and specific information from the data source, the fact that MOCSS model aligns the shared information and imposed the orthogonal penalty during the post-processing process may loss some important information.

In this study, we propose a novel architecture of AE model for multi-omics data integration, where the joint component is derived from the concatenated data sources and the individual component come from the corresponding individual data source. To encourage the model to separate and extract the joint/shared information contained between different omic data and the specific information contained in each data source, an additional orthogonal penalty is applied between the joint and the individual embedding layers. For comprehensiveness, 3 different version of orthogonal penalty was defined and compared, and the model without orthogonal penalty was also compared. Unlike MOCSS model, the joint component and orthogonal penalty are natural components of our model structure. For comparison, we implemented aforementioned AE structures and compared them on a comprehensive simulation setting in terms of input reconstruction and classification accuracy. For further validation, we also applied them on a real-world cancer dataset from TCGA and compared their performance with a dimension reduction model: Joint and Individual Variation Explained (JIVE), which is based on PCA [4]. We chose to compare with JIVE because it similarly imposes an orthogonality constraint between the joint and individual components. It should be noted that, for a fair comparison, all AE models implemented in this study do not contain variational components.

## Methods

### Construction of AE models of different architectures

To compare the effectiveness of multi-omics data integration methods, different AE models were first applied to simulation dataset and further validated on the real-world cancer dataset. We implemented the previously proposed AE models for omics data integration, CNC AE, X AE, MM AE and MOCSS, as shown in Fig. 1a-d. The proposed novel AE structure, Joint and Individual Simultaneous Autoencoder (JISAE) is shown in Fig. 1e, where the encoder starts with the individual omics data as well as their concatenation. These three inputs were separately processed by 4 fully connected layers including the separate final embedding layers for three inputs. To encourage the model to separate and extract the joint (shared) information between the 2 different omics data sources and the individual (specific) information unique to each data source, in addition to the reconstruction loss, an additional orthogonal penalty is applied between the embedding layer of joint and the individual inputs as shown in the orthogonal loss diagram in Fig. 1e. We refer to the models with an orthogonality penalty as JISAE-O. To evaluate the importance of the orthogonality penalties for the model performance, the JISAE model without orthogonal constraints is implemented for comparison, as shown in Fig. 1f.

**Fig. 1.**
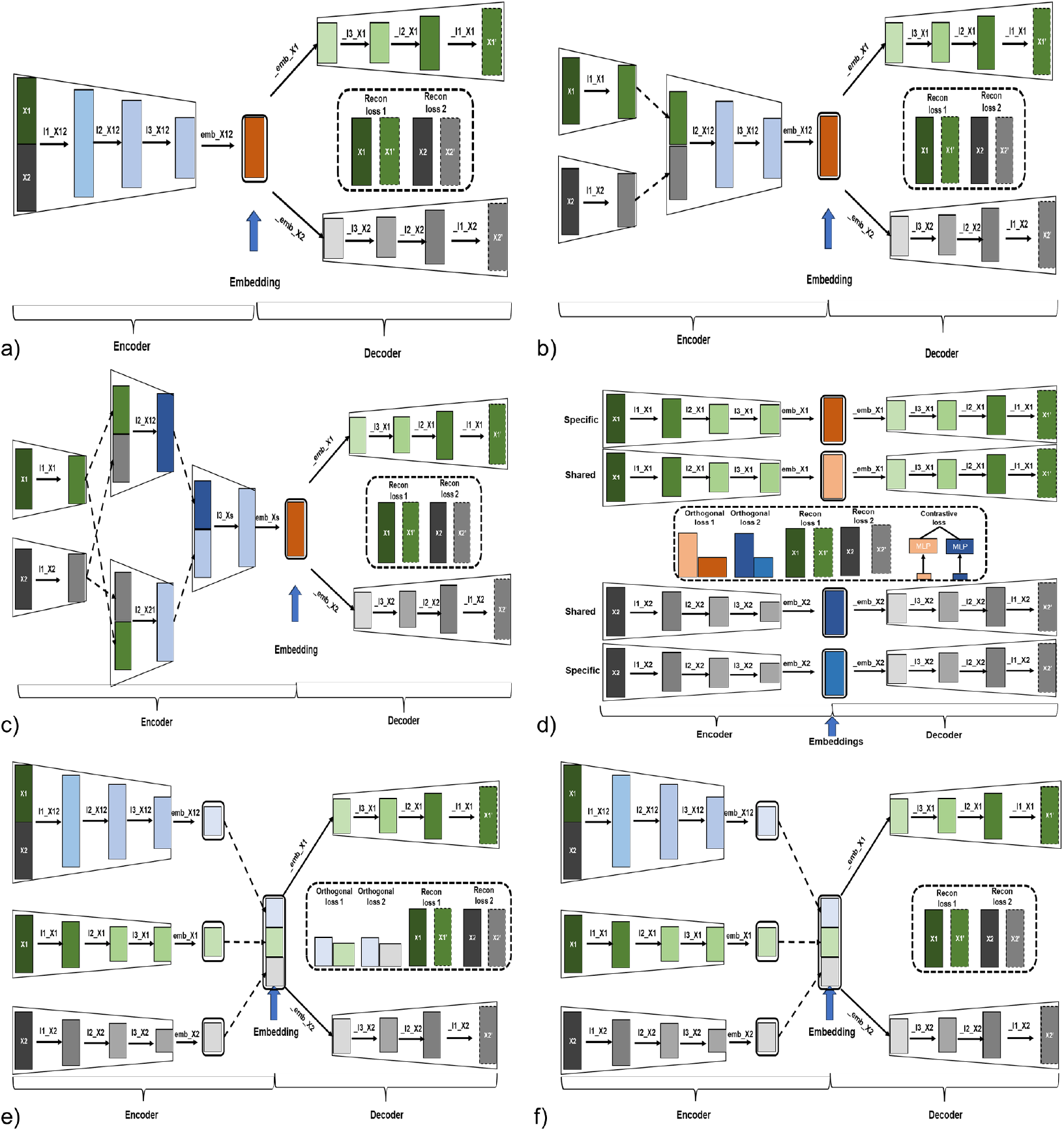
Different AE model structures compared in this study. a): AE with direct X1, X2 concatenated together as the model input, referred as CNC AE; b): AE with processed X1, X2 concatenated together as the model input, referred as X AE; c): AE with different concatenation of processed X1, X2 as the model input, referred as MM AE; d): AE structure from previous study with orthogonal constraints imposed during the post-processing, referred as MOCSS [2]; e): AE with both individual and concatenated X1, X2 as the model input with additional imposed orthogonal constraints, referred as JISAE-O; f): similarly to model in e) without additional orthogonal constraints, referred as JISAE.

To estimate the parameters for each of the AE models, the MSE between the inputs and the reconstructed inputs was used as the loss function for the optimization algorithm. Let 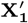 and [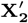 e the reconstructed inputs for **X**_**1**_ and **X**_**2**_ respectively from AE models other than MOCSS (All of these matrices are normalized) and the reconstruction loss can be expressed in (1):

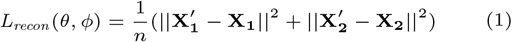

For the MOCSS model, for each data source, there are two reconstructed inputs from shared and specific AE models as shown in Fig. 1 f. Let 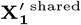 and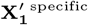 represent the reconstructed inputs from shared and specific AE models respectively for **X**_**1**_ and let 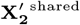 and 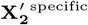 represent the reconstructed inputs from shared and specific AE models respectively for **X**_**2**_ (All of these matrix are normalized). The reconstruction loss for the MOCSS model can be expressed in (2):

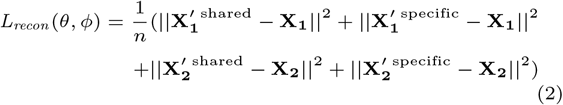

In the MOCSS model, the following penalty was imposed between the shared and specific embedding for each data source. Let **A**_**1**_^shared^, and **A**_**1**_^specific^ to represent the shared and specific embedding for **X**_**1**_, and **A**_**2**_^shared^ **A**_**2**_^specific^ to represent the shared and specific embedding for **X**_**2**_ as shown in Fig. 1f (All of these matrix are normalized). Then the orthogonal loss is calculated as shown in (3), where the ⊙ represents the element-wise product and avg represents the average.

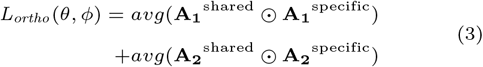

To compare the effectiveness of different orthogonal loss penalties in our proposed AE structure, 3 different orthogonal loss functions were implemented resulting in three different JISAE-O models (distinguished as JISAE-O1, JISAE-O2, and JISAE-O3). Specifically, let **A**_**12**_^shared^ to represent the embedding from joint input **X**_**1**_ and **X**_**2**_, **A**_**1**_^specific^ and **A**_**2**_^specific^ to represent the embedding from individual input **X**_**1**_ and **X**_**2**_ respectively (All of these matrix are normalized). The orthogonal loss for JISAE-O1 that is similar to that of the MOCSS model and is defined in (4).

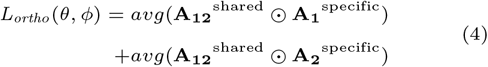

With the same notation for embedding, the orthogonal loss for JISAE-O2 and JISAE-O3 is defined in (5) and (6) respectively:

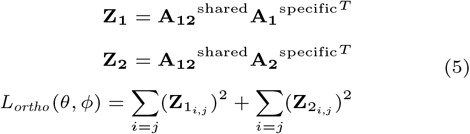

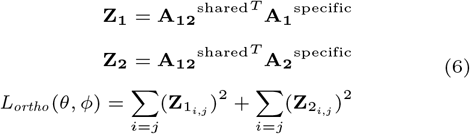

The rows in **A**_**12**_^shared^, **A**_**1**_^specific^, and **A**_**2**_^specific^represent individual subjects. The columns from each matrix represent elements of features from the low dimensional representation for the shared and individual input respectively. Therefore the orthogonal loss defined for JISAE-O2 (see equation 5) imposes the orthogonality constraints among shared and specific embedding for each subject whereas the orthogonal loss in JISAE-O3 (see equation 6) imposes the orthogonality constraints among shared and specific embedding across all subjects in a given batch of training examples for a particular elements of the features. Therefore, the total loss for the MOCSS and JISAE-O models is defined as *L*_*total*_ = *L*_*recon*_ + *λ* * *L*_*ortho*_. *λ* is a hyperparameter that controls the relative contribution of the orthogonal loss to the total loss in these models. The total loss for other AE models is simply *L*_*total*_ = *L*_*recon*_. With the total loss function defined above, the gradient-based optimization algorithm was used to reduce the loss by iteratively optimizing the model weights.

The R package r.JIVE [6] was used to compare the AE methods with JIVE. All AE models in this study were created using the Pytorch framework (1.13.1) [7] with CUDA Toolkit 11.4. The KFold from the sklearn (1.0.2) [9] was used to split the dataset randomly for cross-validation (CV). The Optuna (https://github.com/optuna/optuna) was used for hyperparameter searching based on the average reconstruction loss on the validation set during the 5-fold cross-validation.

### Dataset generation

To compare the effectiveness of multi-omics data integration with different models, we benchmark different AE performance first on a simulation dataset generated by using R package MOSim [5] and further validated on a dataset based on TCGA. For the simulation, RNA-seq (with a total 20,531 features) and miRNA-seq (with a total 1,046 features) were jointly generated from a series of settings. Specifically, data of 2, 3, 4, and 5 groups were generated separately and for each group setting, 20%, 40%, 60%, 80% and 100% of the genes were set to be differentially expressed. Thus a total of 20 different dataset were simulated representing different level of difficulty to test the model and the total number of samples were kept to 600 in each dataset as shown in Table 1.

**Table 1.** Simulation dataset generated from 20 different settings with increasing level of difficulties from top to bottom, right to left used for model training and testing in this study. (% DEG: % of differentially expressed genes)

| % DEG | 20% | 40% | 60% | 80% | 100% |
| --- | --- | --- | --- | --- | --- |
| Groups |  |  |  |  |  |
| 2 | 300 samples/group | 300 samples/group | 300 samples/group | 300 samples/group | 300 samples/group |
| 3 | 200 samples/group | 200 samples/group | 200 samples/group | 200 samples/group | 200 samples/group |
| 4 | 150 samples/group | 150 samples/group | 150 samples/group | 150 samples/group | 150 samples/group |
| 5 | 120 samples/group | 120 samples/group | 120 samples/group | 120 samples/group | 120 samples/group |

For the cancer dataset, multi-omics TCGA preprocessed datasets were downloaded from https://acgt.cs.tau.ac.il/multi_omic_benchmark/download.html [11], which includes gene expression, and miRNA expression data as well as patient clinical information of 10 different cancers (AML: Acute Myeloid Leukemia, BIC: Breast cancer, COAD: Colon adenocarcinoma, GBM: Glioblastoma multiforme, KIRC: Kidney renal clear cell carcinoma, LIHC: Liver hepatocellular carcinoma, LUSC: Lung squamous cell carcinoma, SKCM: Skin Cutaneous Melanoma, OV: Ovarian serous cystadenocarcinoma and SARC: Sarcoma). Not all subjects have both omic measurements, therefore, for a given cancer type, we only kept the subjects that have both omic data. For a given omic measurement, the number of measured features can be different across the cancer types, we selected the cancer types that have the most consistent feature dimensions across both omic measurements. After this filtering, 6 cancer types including BIC, LUSC, SKCM, LIHC, SARC, and KIRC were left for subsequent analysis.

### Hyperparameter tuning for AE models

To choose the optimal hyperparameters for different AE models on the simulation datasets, with each dataset, 80% were allocated for training while the remaining 20% were used for final model evaluation as shown in Fig. 3a. The 5-fold cross-validation (CV) was used on the training set to choose the optimal hyperparameters for each AE model corresponding to the model configuration with the lowest average reconstruction loss on the validation set. It is computationally impossible to cover the entire hyperparameter space. Therefore, Optuna https://github.com/optuna/optuna a hyperparameter optimization framework for hyperparameter searching was used by conducting 50 random trails for each model in the simulation setting and 100 random trials for cancer dataset, where each trial corresponds to a different combination of hyperparameters. The selected optimal hyperparameters were then used to retrain the model on the entire training set in order to extract the embeddings of interest for both training and testing set for subsequent analysis. For the cancer dataset, after excluding solid tissue normal and metastatic samples as shown in Fig. 2a, the primary tumor samples were divided into non-overlapping training and testing sets following the same partition rule as shown in Fig. 2b with the same training strategy shown in Fig. 3a.

**Fig. 2.**
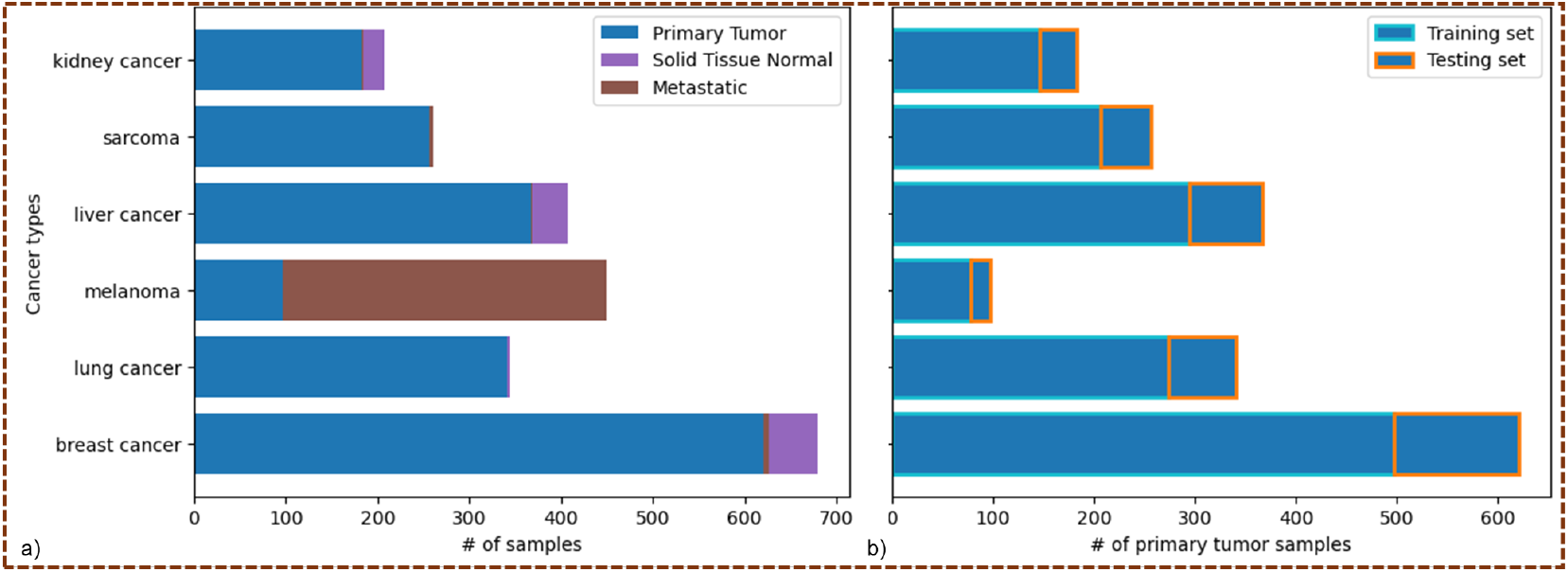
TCGA dataset used for model training and testing in this study. a): TCGA samples downloaded from https://acgt.cs.tau.ac.il/multi_omic_benchmark/download.html with gene and miRNA expression data available; b): Primary tumor samples were used for training and testing.

**Fig. 3.**
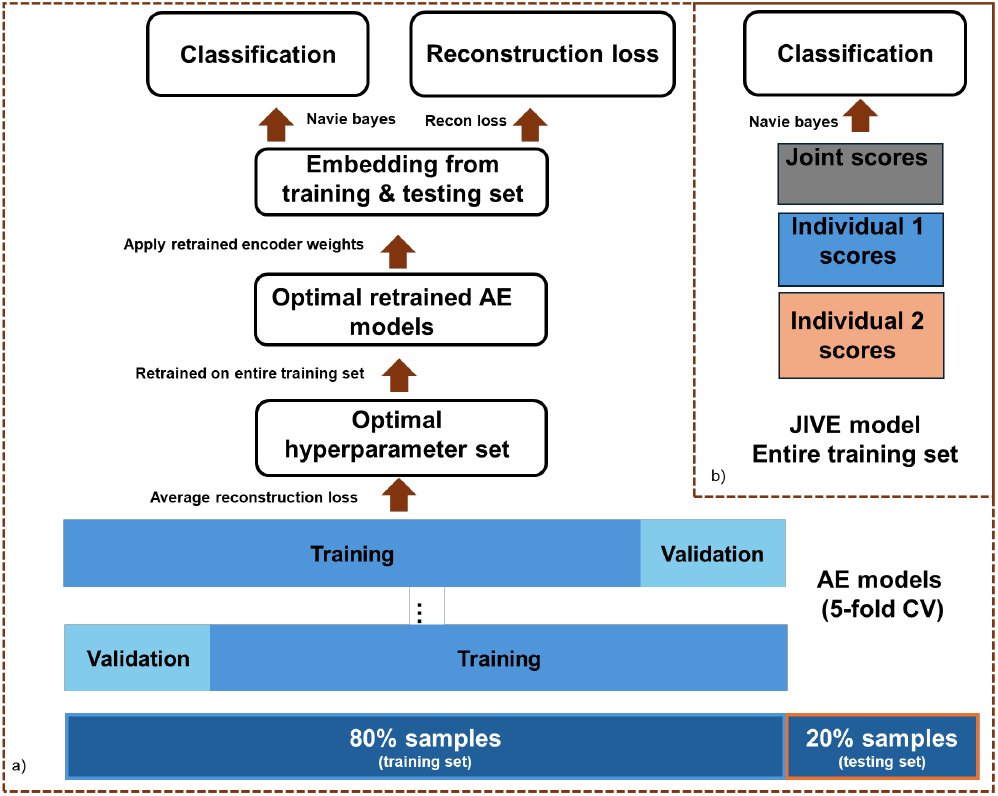
Model training and comparison process in this study.

With the cancer dataset we also compared the performance between AE models and JIVE. To extract the joint components across gene and miRNA expression data and individual components that are specific for gene and miRNA expression data respectively defined in the JIVE model, we applied the JIVE function implemented in r.JIVE package to the entire training set with the default parameter settings (see Fig. 3b). Specifically, the gene expression and miRNA expression training set were put in a linked list and the JIVE algorithm was then used to find the low-rank approximation matrix for the joint and individual components respectively with the permutation test to find the optimal ranks. After obtaining the low-rank approximation of joint and individual matrix, with the given rank, the prcomp function from the R stats package [10] was then used to decompose these low-rank matrices as the product of the loading and the corresponding score matrix. The score matrices for the joint and individual components were then concatenated together as the new features for the subsequent analysis as shown in Fig. 3b. With the loadings calculated based on the training set, the jive.predict function implemented in r.JIVE was then used to obtain the score matrices for the joint and individual components on the testing set. The joint and individual components were similarly concatenated together for subsequent analysis.

In terms of hyperparameter selection choices for AE models, the number of nodes for all the layers (except the first and last layer determined by the dimensions of the inputs) across all AE models are chosen from {32, 64, 128, 256, 512, 1, 024}. Compared with the MOCSS model, batch normalization followed by dropout layers was included in all other AE models, and the dropout rate was selected from {0, 0.1, 0.2, 0.4, 0.6}. In terms of model structure, whereas all other AE models can have different numbers of nodes in each layer (the dimensions for the shared and specific embedding for JISAE-O and JISAE models were kept the same), the MOCSS model is symmetric where the number of nodes across the encoder mirrors the number of the nodes across the decoder. In terms of nonlinear activation functions, instead of ReLu, hyperbolic tangent activation is used in the MOCSS model. In terms of hyperparameters for model training, the learning rate is chosen from 1*e*^−5^ to 1*e*^−2^ on a logarithmic scale; l2 penalty for model weights is chosen from 1*e*^−8^ to 1*e*^−5^ on a logarithmic scale; number of training epoch is chosen from {30, 60, 90, 120, 150}; batch size was chosen from {32, 64, 128, 256, 512} and the Adam optimizer was used for model weights optimization. Finally, for the three AE models with orthogonal loss constraints, as mentioned above, *λ* was chosen from {10^−3^, 10^−2^, 10^−1^, 10^0^, 10^1^10^2^, 10^3^} to control the relative contribution of the orthogonal loss to the total loss.

### Performance comparison of different models

To assess whether the AE models can compress the data and retain the essential information for the data, reconstruction loss on the entire training and testing set was calculated by following (1) and (2) for AE and MOCSS models respectively after applying the optimal retrained AE models to training and testing set separately as depicted in Fig. 3a. To investigate whether the omic data integration via AE can be useful for subsequent analysis, a Gaussian naive Bayes classifier from the the sklern package [8] that doesn’t require any additional hyper-parameters was used for the classification of different groups with the extracted embeddings. This comparison was conducted for both simulation and cancer dataset. For the cancer dataset, we additionally compared the AE model with JIVE. Essentially, the concatenated joint and individual components extracted from the JIVE model were also used for classification. To assess the effectiveness of AE and JIVE models, the original given features as well as their concatenation were also used as the baseline. Specifically, the gene expression and miRNA expression data in the training and testing set were normalized by using the MinMaxScaler from the sklearn package [8] separately. MinMaxScaler scaled the data set such that all feature values are in the range of [0, 1]. For a fair comparison, the input for all AE and JIVE models was also scaled by MinMaxScaler for training and testing sets separately.

## Results

### Simulated data

Data sets with different level of difficulties were simulated from 20 settings. 600 samples were simulated from each setting, with 2, 3, 4, or 5 groups containing 300, 200, 150 or 120 samples respectively, as shown in Table 1. The total number of features for gene and miRNA expression in the simulation data matched the total number of features in original TCGA dataset. The TCGA dataset contained 685 BIC, 221 COAD, 344 LUSC, 291 OV, 450 SKCM, 410 LIHC, 261 SARC, 170 AML, 274 GBM, and 208 KIRC subjects that had both omics measurements. In the gene expression data across these 10 cancer types, GBM contained expression measurements for a total of 12,042 genes while the rest of the cancer types all contained the expression level measurements of 20,531 genes. In the miRNA expression data, six cancer types, BIC, LUSC, SKCM, LIHC, SARC, and KIRC, had expression measurements of 1,046 miRNAs. Three cancer types, COAD, OV, and AML, had the expression measurements of 705 miRNAs, and GBM contained expression measurements of 534 miRNAs. To make sure each omics data had the same number of features across the selected cancer types, 6 cancer types including BIC, LUSC, SKCM, LIHC, SARC, and KIRC were selected for subsequent analysis. Some of these cancer types include multiple sample types (*e*.*g*., primary tumor, solid tissue normal, and metastatic samples), but we restricted our analysis to primary tumors only (*i*.*e*., a total of 1,866 subjects) with 621, 341, 97, 367, 257, and 183 for breast, lung, melanoma, liver, sarcoma and kidney tumor samples respectively, as shown in Fig. 2a.

### Selected hyperparameters for AE models

The optimal hyperparameters for each AE model were selected based on the lowest average reconstruction loss on the validation set during the 5-fold CV among Optuna trials. The lowest average 5-fold CV reconstruction loss among different AE models for each simulation dataset is depicted in Fig. 4 and the corresponding optimal hyperparameter set for each AE model is shown in Supplementary Table 1. Clearly, the average reconstruction loss are increasing as the simulation dataset become difficult (see Table 1). The lowest average 5-fold CV reconstruction loss on cancer dataset is shown in Fig. 5 and supplementary Table 2 depicted the corresponding optimal hyperparameter set. It should be noted that the reconstruction loss for MOCSS model were omitted, since it includes two separate reconstruction losses for each data source (from shared and specific AE models as shown in Fig. 1d also see the reconstruction loss for the MOCSS model in (2)) making the comparison incompatible.

**Table 2.** Reconstruction loss comparison of TCGA dataset based on the retrained model weights on the entire training set.

| Features | Embedding dimension | Reconstruction loss <sup>1</sup> | Reconstruction loss <sup>2</sup> |
| --- | --- | --- | --- |
| embedding (CNC_AE) | 32 | 0.0306 | 0.0737 |
| embedding (X_AE) | 256 | 0.0303 | 0.0729 |
| embedding (MM_AE) | 512 | 0.03 | 0.0757 |
| embedding (JISAE) | 1,536 | 0.0276 | 0.072 |
| embedding (JISAE-O1) | 3,072 | 0.0255 | 0.0795 |
| embedding (JISAE-O2) | 3,072 | 0.0292 | 0.0838 |
| embedding (JISAE-O3) | 192 | 0.0312 | 0.0754 |

<sup>1</sup>Calculation based on the training set that contains n=1494 subjects; <sup>2</sup> Calculation based on the testing set that contains n=372 subjects;

**Fig. 4.**
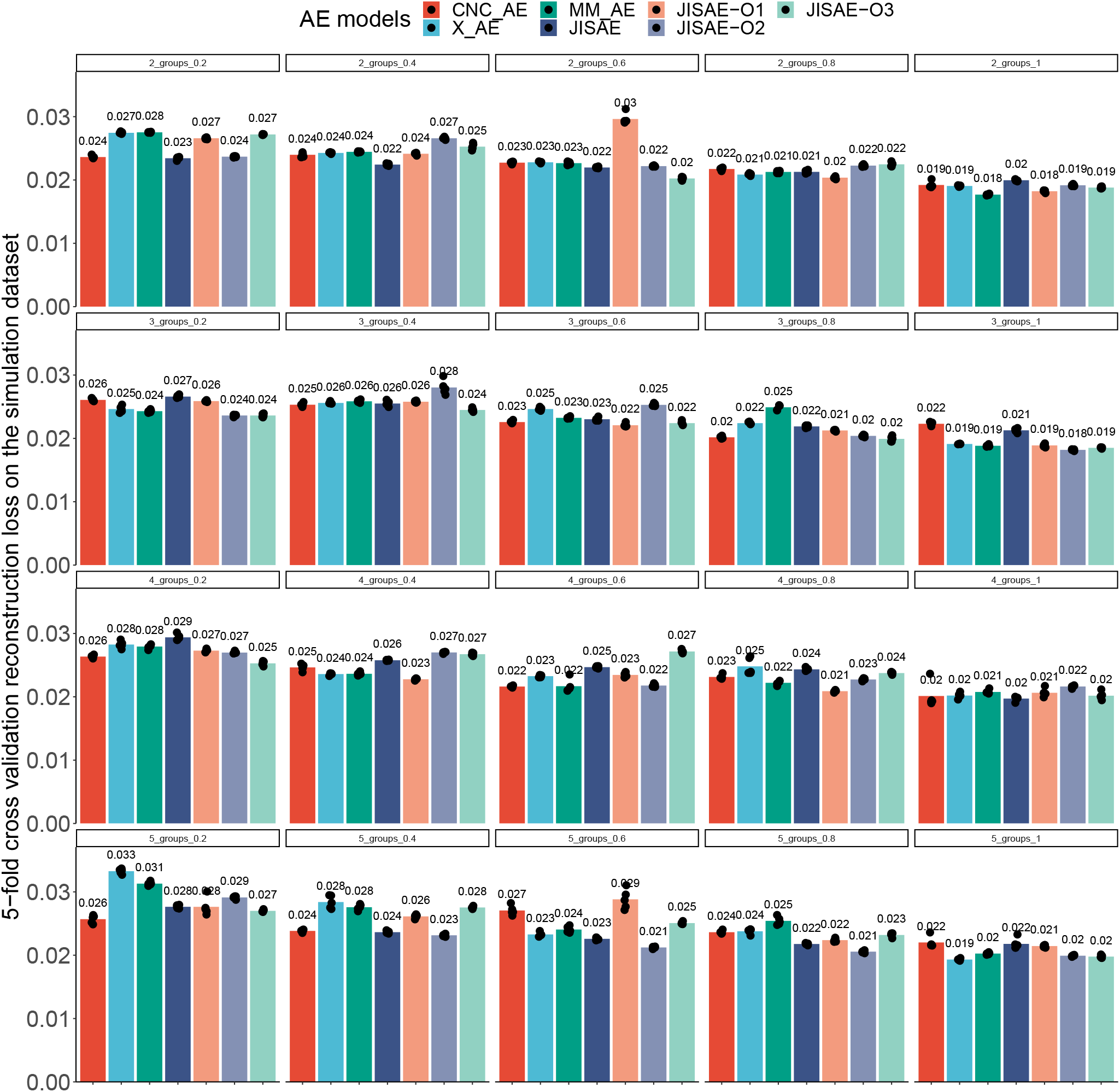
The lowest average 5-fold CV reconstruction loss among different AE models for each simulation dataset.

**Fig. 5.**
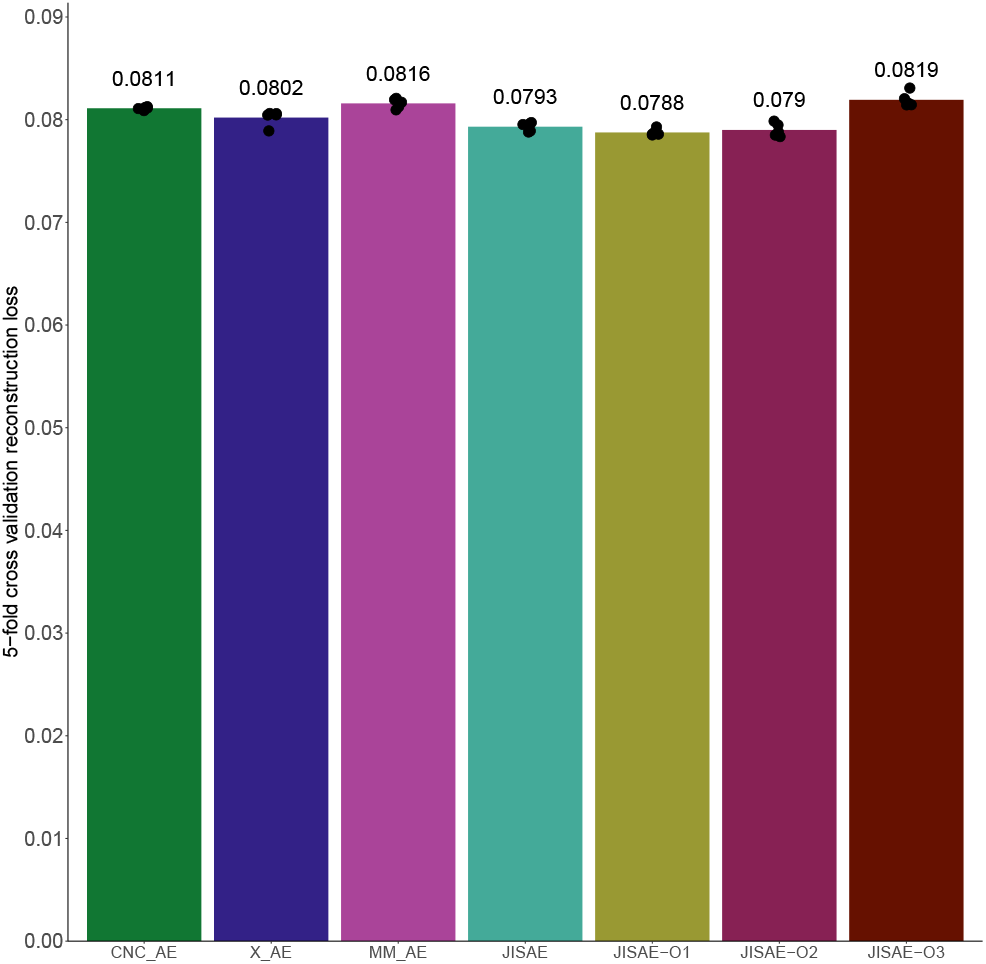
The lowest average 5-fold CV reconstruction loss among different AE models for TCGA cancer dataset.

### Comparison of model performance

The optimized hyperparameters based on the lowest average reconstruction loss were used to retrain all the AE models on the entire training set separately. To evaluate the generalization of AE models for data reconstruction, the average reconstruction loss on the training and testing set was calculated by following (1) for all AE models other than MOCSS based on the retrained model weights. The averaged reconstruction loss on the simulation training and testing dataset is shown in Fig. 6a and 6b respectively. As expected, with the increase of the difficulty across the simulation dataset, the averaged reconstruction loss were in general increased on both the training and testing set indicated by the yellow to red color in Fig. 6. The extracted embeddings were used as the new features in a Gaussian naive Bayes classifier for classification task with 5-fold CV. As shown in Fig. 6c and 6d for the simulation training and testing dataset respectively, almost all the data points in both datasets were classified with accuracy of 100%.

**Fig. 6.**
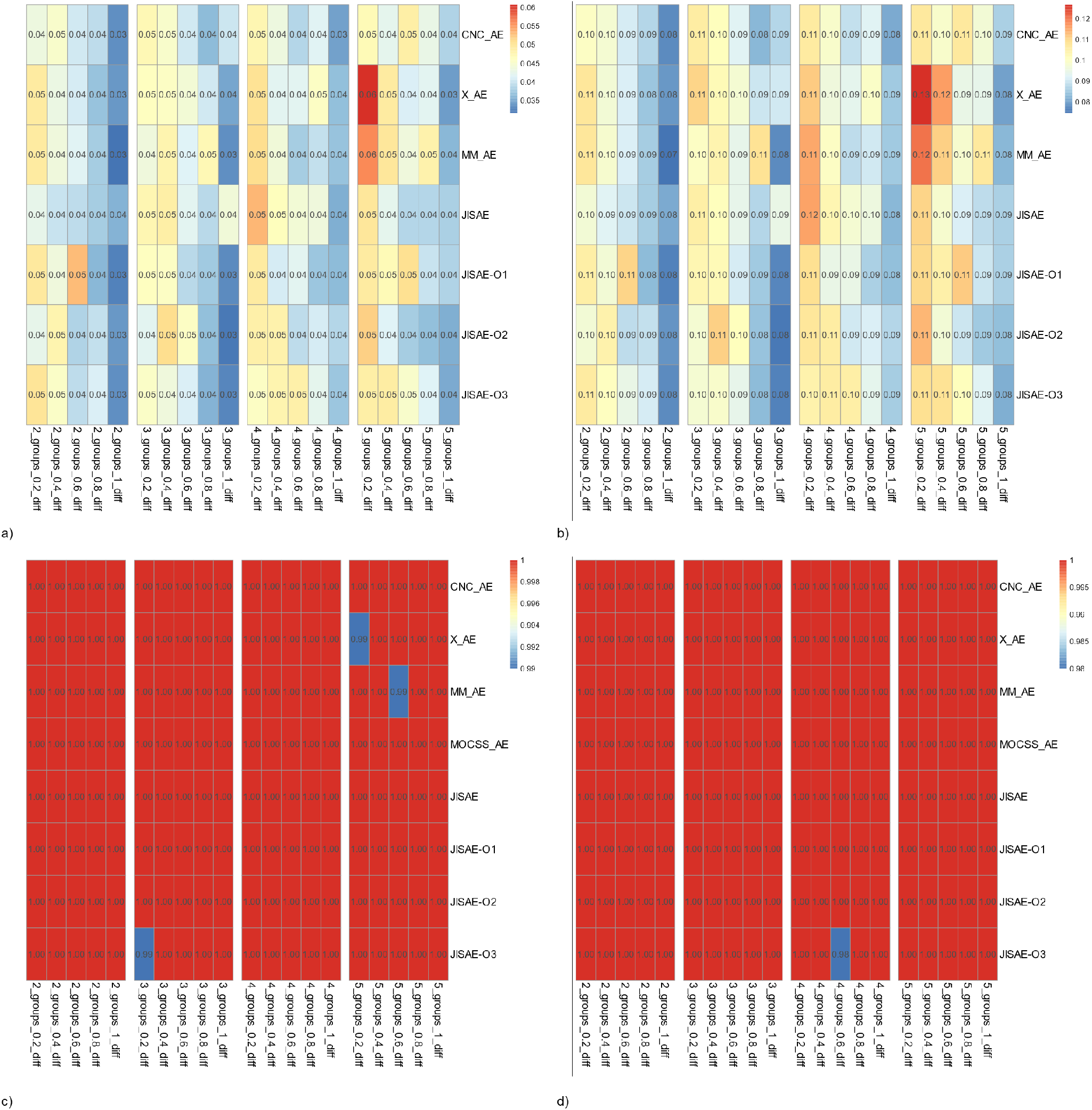
Average reconstruction loss and classification accuracy for AE models on each of the simulation dataset. a): reconstruction loss on training dataset; b): reconstruction loss on testing dataset; c): classification accuracy on training dataset; d): classification accuracy on testing dataset.

For the TCGA cancer dataset, the optimal dimensions of embeddings and corresponding reconstruction loss for each AE model except MOCSS were included in Table 2. The dimension of embedding ranges from 32 with the CNC AE model to 3,072 for JISAE-O1 and JISAE-O2 models. Therefore in terms of omic data reduction, CNC AE reduced the dimension the most from 21,577 (including 2,0531 gene expression and 1,046 miRNA expression) to 32 followed by JISAE-O3 with 192 dimensions for their embeddings. In terms of the reconstruction loss, on the training set, JISAE and JISAE-O1 achieved relatively lower reconstruction loss compared to all the other AE models. On the testing set, CNC AE, X AE, and JISAE achieved relatively lower reconstruction loss in comparison.

The model performance was also compared with JIVE, which decomposes the original concatenated matrix into the joint matrix denoted as **J** with separated individual matrices denoted as **A**_**1**_ and **A**_**2**_ for gene expression and miRNA expression respectively in addition to a residual noise matrix denoted as **R** [4]. Using the permutation test in the r.JIVE package [6], we selected a rank of 6 for the joint matrix and ranks of 160 and 100 for individual matrices (*i*.*e*., **A**_**1**_ and **A**_**2**_) respectively. The score matrices for joint and individual components were then extracted with PCA as mentioned earlier. On the testing set, by directly applying the loadings from the training set to the testing set, we obtained the corresponding joint and individual score matrices with the same ranks as the training set.

To evaluate the effectiveness of the embeddings from the AE models and the joint and individual components from JIVE model on the TCGA cancer dataset in the subsequent analysis, the Gaussian naive Bayes classifier was used for cancer type classification. We performed a 5-fold CV and reported the average classification accuracy on both the training and testing set using the corresponding embeddings extracted from retrained AE models as illustrated in Fig. 3a. As shown in Fig. 3b, the joint and individual components from the JIVE model were concatenated together for cancer classifications. As a baseline, the original features were also used for comparison.

On the training set, as shown in Fig. 7a, in general, using original features produced a lower classification accuracy, and embeddings extracted from AE models can significantly boost the classification accuracy. With the combined joint and individual components extracted from the JIVE model, the naive Bayes classifier results in the lowest average classification accuracy at 67.2%. With the gene and miRNA expression, we obtained similar classification a ccuracy (both a veraged about 79%), and the direct concatenation of the two surprisingly further reduced the accuracy to 71.42%. Among all the AE models, the embeddings from MM AE produced the highest classification a ccuracy o f 9 5.18% f ollowed b y J ISAE (93.51%), JISAE-O1 (92.77%) and JISAE-O2 (91.68%), while embeddings from JISAE-O3 produced lowest accuracy of 80.45% among all tested AE model architectures. For each AE model, by applying the trained model weights from the entire training set to the testing set, we obtained the embeddings for the testing set. The classification a ccuracy o n t he t esting s et i s s hown i n Fig. 7b. The original features generated much lower classification accuracy (all around 58%) compared with AE and JIVE models.

**Fig. 7.**
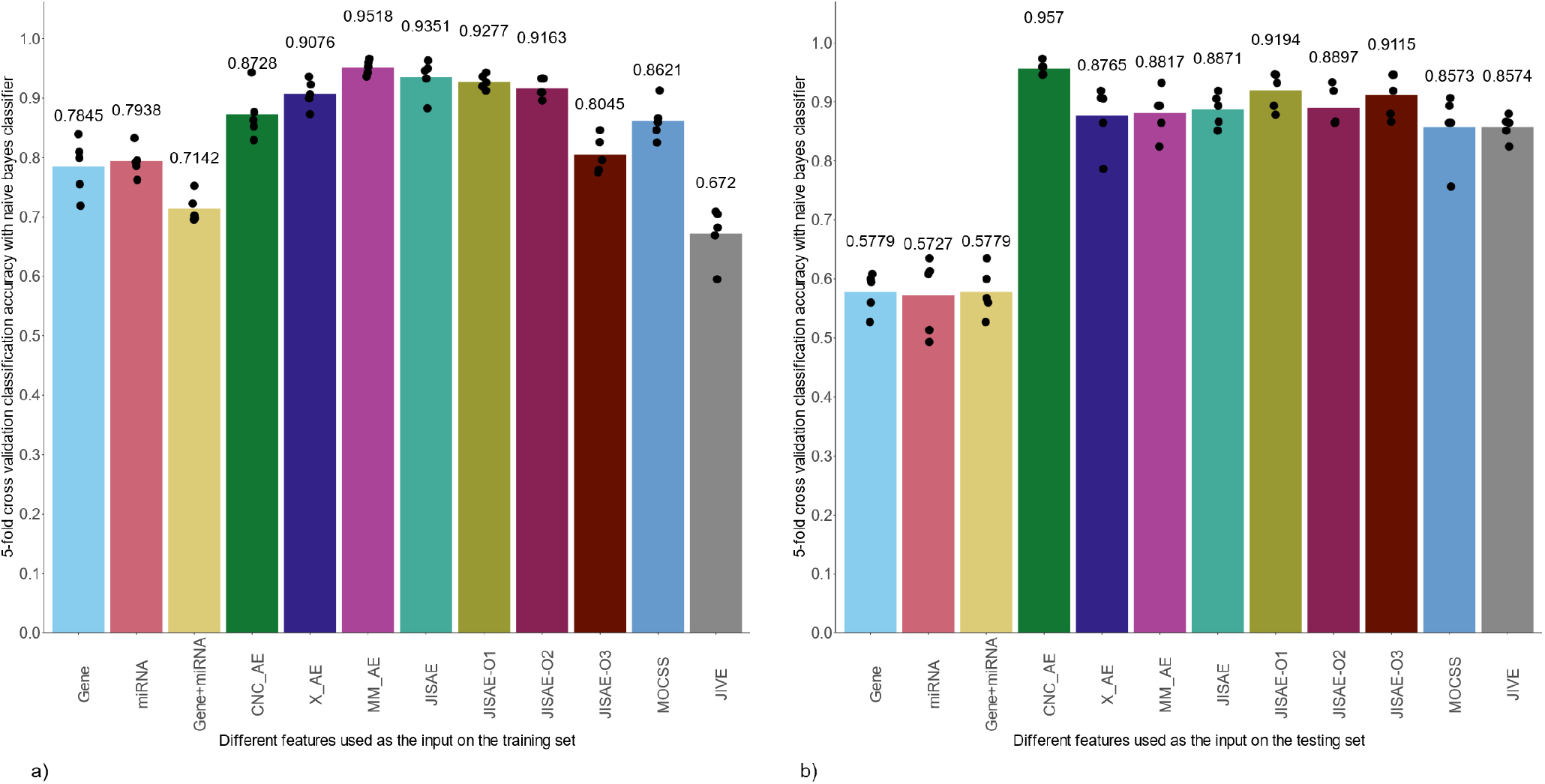
Average 5-fold CV accuracy using different features as input for naive Bayes classifier to classify 6 different types of cancers. a) on the training set; b) on the testing set

While the CNC AE model achieved 87% average classification accuracy on the training set, it achieved the highest average accuracy of 95.7% on the testing set. In comparison, JISAE-O1 and JISAE-O3 achieved similar performance (around 91%) on the testing sets. Unexpectedly, using joint and individual components from the JIVE model, a much higher classification accuracy of 85.74% was achieved compared to the lowest accuracy of 67.2% on the training set.

## Discussion

In this study, we proposed a flexible new AE model structure for data integration in multi-omics data. Compared to previous integrative AE models, the proposed models use both shared and specific inputs and allow flexibility to incorporate orthogonality in the loss function. We compared the reconstruction loss and classification accuracy of the novel model structures to four existing integrative autoencoders: CNC AE, X AE, MM AE, and MOCSS [12, 2].

To train the AE models and compare the model performance, simulated datasets with increasing levels of difficulties were used for reconstruction loss (except MOCSS) as well as classification accuracy comparison. To evaluate the model on the real-world dataset, the pre-processed gene expression and miRNA expression data of 6 different cancer types from TCGA dataset were used [11]. The JIVE model was also evaluated on the TCGA dataset for comparison. In both settings, the whole dataset was split into the training and testing set to train the model and evaluate the model generalization error respectively as shown in Fig. 3. For the AE models, the objective is to reconstruct the input with the learned lower dimensional embedding. Therefore, the 5-fold CV on the training set was used to select the optimal hyperparameter set that corresponds to the lowest average reconstruction loss on the validation set (see Fig. 4 and Fig. 5 for reconstruction loss on simulation and TCGA datasets respectively). The AE models were then retrained on the entire training set with the fixed optimal hyperparameter selected earlier and used to compare the reconstruction loss (both for the training and testing set), the corresponding embeddings were used for the subsequent classification task.

In terms of model performance on the simulation datasets, as shown in Fig. 6a and b, reconstruction loss are in general getting smaller as the dataset become easier (*i*.*e*., from left to right within each group) on both training and testing set respectively. On the most difficult dataset that contains 5 groups with 20% differentially expressed genes (DEGs), X AE and MM AE showed slightly higher reconstruction loss on both training and testing set compared with other AE models. Despite the slightly difference in reconstruction loss among different AE models, the embeddings extracted from AE models are all seems informative in terms of classification of different groups (see Fig. 6c and d).

On the TCGA training set, JISAE-O1 achieved the lowest average reconstruction loss among all AE models as shown in Table 1, indicating the effectiveness of this AE model in reconstructing the model input with the learned embeddings. Although the reconstruction loss is much higher on the testing set compared with the training set, the JISAE model achieved the lowest reconstruction loss (0.072) on the TCGA testing set, indicating its better generalizing ability to reconstruct the input for the unseen data. In terms of learned embeddings, CNC AE reduces the dimension of the input the most, with 32 new features to represent the original 21,577 features. Overall, the different AE architectures constructed in this study showed slightly different capacities to capture the essential input information and generalize that to the unseen data.

These learned embeddings from AE models were then used for subsequent cancer classifications. For the JIVE model, the joint and individual components based on the entire training set were extracted and combined for the subsequent classification. The loadings learned on the training set were used to compute the joint and individual components for the testing set followed by classifications as well. As shown in Fig. 7, embeddings learned in AE models can significantly boost the model performance both on training and testing set respectively, demonstrating the effectiveness of AE models to extract the low dimensional information from the multi-omics data source for subsequent data analysis. With joint and individual components from the JIVE model, although the naive Bayes classifier results in the lowest average classification accuracy at 67.2% on the training set, a much higher classification accuracy of 85.74% was achieved on the testing set. This performance is on par with the MOCSS model introduced in a previous study [2].

Comparing the model performance in terms of reconstruction loss derived from the embeddings and subsequent classification tasks with these embeddings, it is clear that the low reconstruction loss is not directly translatable to subsequent higher model performance. For instance, CNC AE is a simple structure, that has the lowest dimension of embeddings and is performing the best for testing set classification, nevertheless, the reconstruction loss for CNC AE is not the lowest one among all AE models constructed in this study. Although the highest average accuracy of 95.7% on the testing set was achieved by the CNC AE model, JISAE-O1 achieved consistently high model performance on both training (92.77%) and testing set (91.94%) indicating the greater robustness of this model. Although not as highly accurate as the JISAE-O1, the MOCSS model also demonstrated consistently higher model performance between the training and testing set (86.21% on the training set and 85.73% on the testing set). The fact that the same orthogonal loss function was used in JISAE-O1 and MOCSS models (see equations (3) and (4) respectively) indicates that this orthogonal loss might be the optimal one compared to the other 2 orthogonal loss (see equations (5) and (6)) defined in this study for the classification tasks.

In theory, deep-learning-based multi-omics data integration methods such as AEs can have more flexible architectures with the possibility of designing specific structures for processing each data source. This flexibility and relatively higher model performance in this study demonstrated its potential superiority over the JIVE model for data integration. Although all AE models considered in this study achieved similarly high model performance on the simulation dataset in terms of reconstruction loss and classification accuracy, with the real-world TCGA dataset, the novel AE model structure coupled with orthogonal penalties proposed in this study showed relatively higher model performance compared with previous MOCSS model (see Fig. 7b).

However, there are several limitations to the current study. First of all, although 5-fold CV was used to select the optimal hyperparameter for each AE model based on the average reconstruction loss on the validation set, L2 loss and dropout layers were also implemented in the models, a relatively large discrepancy still appears between the classification accuracy on the training and testing set for some of the AE models with TCGA dataset. To mitigate this issue, more data could be collected for model training and evaluation in the future, and variational AE models that contain intrinsic model regularization can be used to replace the AE structures in this study. Secondly, we only evaluated the model performance with the naive Bayes classifier, which could potentially bias our comparison results if the embeddings from AE models or joint and individual components from the JIVE model do not uniformly satisfy the underlying model assumption (*i*.*e*., the conditional independence of features given a class label). Therefore, more comprehensive evaluation metrics derived from different machine learning models can be used in the future for a more objective evaluation. Finally, the AE models were trained to reconstruct the model input, therefore, as mentioned earlier, the embeddings that can be effectively used to reconstruct the model input might not directly benefit the subsequent analysis (*e*.*g*., classification in this study). However, for a fair comparison with JIVE, which is not given any labeling information, reducing the reconstruction loss on the validation set during the model training is a reasonable loss to minimize. For future studies with AEs, in addition to the reconstruction loss, the customized loss function that is directly related to the specific tasks can be added to the model training process (*e*.*g*., classification error), such that the extracted embeddings not only contain essential information in the input data but are also directed towards better performance for a given specific subsequent analysis when these embeddings are used as the new features for the analysis.

## Data Availability Statement

The preprocessed TCGA data was obtained from https://acgt.cs.tau.ac.il/multi_omic_benchmark/download.html [12].

## Competing interests

No competing interest is declared.

## Author contributions statement

C.W. and M.O. conceived the study. C.W. implemented the models and performed the analysis. C.W. and M.O. wrote and reviewed the manuscript.

## Acknowledgments

This work was supported by the Miami University start-up fund (M.O.).

## Notes

https://github.com/wangc90/AE_Data_Integration

### Competing Interest Statement

The authors have declared no competing interest.

